## Supplementary Sections for "Population-specific genome graphs improve high-throughput sequencing data analysis: A case study on the Pan-African genome"

### S1 Graph construction

#### S1.1 Graph Building Pipeline

This pipeline takes a number of input population variants and GRCh38 raw reference and outputs the final linear reference and the graph for downstream tools. Workflow has four main steps (see Figure S1a):

- **ALT processing:** takes GRCh38 linear reference in combination with the ALT contig mappings and produces a variant file for the ALT variants as well as the initial linear reference
- **Prepare inputs:** takes a number of input variants (VCF files) and performs the necessary initial process (filter, split, normalize) for merging.
- **Merge variants:** Combines variants from multiple sources and produces the intermediate graph VCF with required annotations
- **Filter variants:** Performs two filtering steps (Structural Variation filter and Multimap filter) on the intermediate graph VCF and generates the final linear reference and the graph VCF

##### *ALT processing*

This step produces the intermediate linear reference as well as the variations from ALT contigs Intermediate linear reference is a combination of primary chromosomes, unplaced and unlocalized contigs and the additional decoys defined for the reference. ALT variations are derived from alignments of ALT contigs onto the primary sequences

**Linear reference** Linear reference is constructed using primary chromosomes, unplaced and unlocalized contigs from the assembly and decoys (HS38D1 and EBV) defined for the reference. ALT and NOVEL contigs are excluded since they will be represented as variations on the primary chromosomes and FIX contigs are ignored.

**ALT variant generation** ALT variant generation starts with the GFF files provided by GRC. These files describe alignment of ALT contig with respect to primary chromosome in CIGAR format. Preliminary variations are obtained by going through the CIGARs and extracting differences between sequences. In some cases, ALT mapping might contain inverted sequence patches (see Figure S2) and alternative alignments of the subsequence of an ALT are defined for these regions. In these cases, the variants from primary alignment are ignored in these regions and alternative mapping is used instead.

**Variant splitting** Alignments provided by GRC are less greedy and often combine nearby variants into large events even if there are large chunks of identical sequences in between. For example, consider the alignment for chr5_GL339449.2_alt: M29 I623 I2085 I1169 D623 D2085 D1169 M5399... . Consecutive insertion and deletion events can be combined and the alignment can be simplified into M29 I3196 D3196 M53399.... The event I3196 D3196 represents a sequence replacement of 3196 bases from the primary sequence by the ALT sequence. However, the majority of two sequences are identical and this replacement can be reduced into 7 single base changes (SNPs) instead.

In order to obtain a more minimal representation of the variations, these kinds of replacement variations are aligned to each other using Needleman-Wunsch algorithm. If a long sequence of identical matching blocks are identified in the alignment then this replacement variation is split into smaller variations from the matching blocks.

Once all the variations for an ALT contig are obtained, they are left normalized with respect to the linear reference. Finally, variations from all the ALT contigs are merged and outputted as a VCF file.

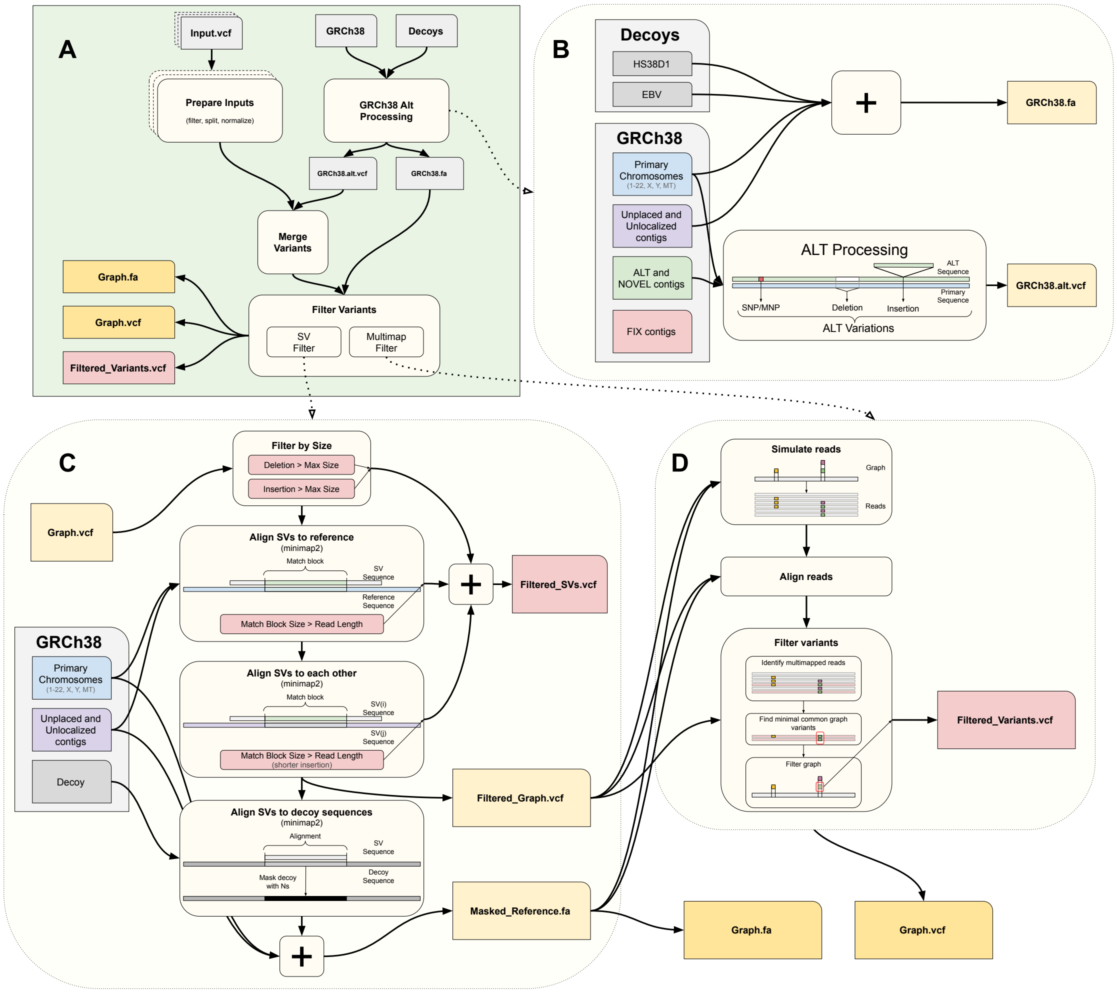


**Figure S1. Graph reference construction pipeline.** The pipeline takes a set of VCF files and the linear reference (to be used as the backbone of the graph reference) as input, and outputs the constructed graph reference along with the modified linear reference and a list of variants that are excluded from the graph. (i) The input variants are processed to avoid any incompatibility with the graph representation and resolve potential issues in the VCF files. (ii) The alt-contigs in the linear assembly are added as edges into the graph reference and decomposed into smaller variants if necessary. (iii) All variants are merged and the allele frequency is re-calculated. (iv) Structural variants that are similar to the linear assembly or to each other are filtered and the decoy sequences are modified. (v) The reads are simulated from the constructed graph to detect and prune edges that cause multi-mapping in read alignment.

##### *Prepare inputs*

Input preparation is a simple step to process VCF files from potentially different sources and unify the variant structure for merging. Steps involve:

- Split multiallelic variants
- Remove non-standard variant definitions and leaving only fully sequence resolved variants
- Apply any additional filtration steps (such as filter by allele frequency) to choose variants to be added

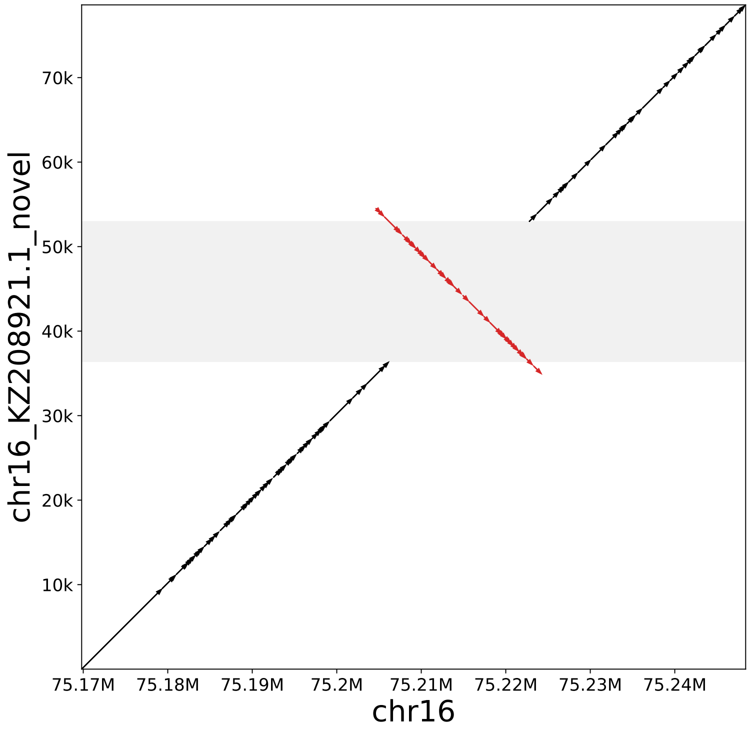


**Figure S2. Dot plot for an alt/novel contig** Black arrows represent similarity in forward strand and red arrows represent similarity in reverse strand. This alt sequence contains an inverted subsequence (gray region) in the middle.

- Left normalize all variants to end up with consistent representation
- Clear unused annotations, ID and FILTER fields and sample information
- Annotate with necessary information to calculate effective allele frequency after merging
- Annotate variants to indicate the original source file using the ID assigned to the VCF file

##### *Merge variants*

Merge variants is a simple step that takes multiple prepared input VCF files and outputs a single biallelic graph candidate VCF file. It combines all variants into a single set while aggregating annotations if the same variant originates from multiple sourcesOnly additional processing is done to calculate the effective allele frequency if the same variation originates from multiple source files. Final allele frequency for a variation is the average of allele frequencies originating from all the source files weighted by the number of samples used for corresponding source file. Produced graph file is then pruned using the variant filtration step to obtain the final graph.

##### *Variant filtration*

Once the source variants are processed and merged, there needs to be some filtration applied in order to obtain a usable graph. Filtration process works in two steps. First, large events (structural variations) are handled in order to eliminate introducing sequences that would result in duplications. Second, remaining smaller variants are scanned for potential multimapping problems and problematic variants are pruned.

**Structural variant filter** Addition of large events (structural variations) may require special consideration for short read analysis. Especially, insertions longer than read size have to go through filtration in order to not make alignments ambiguous. Moreover, implementations of downstream tools can impose maximum size for these events. This filter aims to solve these problems and retain only useful structural variations in the graph. Flow of the process is shown in Figure S1c and details are as follows:

1. Events larger than defined maximum sizes are removed
2. Remaining insertions are aligned to the linear reference (using minimap2). If there is a subsequence (match block) in the insertion that is identical to the non-decoy sequences in the linear reference and is at least read length (default 150bp) size, adding that insertion would introduce ambiguity for some reads. That insertion is filtered.
3. Remaining insertions are aligned to each other (using minimap2). In similar fashion, if there is a common identical subsequence of at least read length, the smaller insertion is filtered.
4. Remaining insertions are added to the graph, but since decoys for a reference are obtained by common additional sequences that are not in the reference, it is possible that some of those sequences are already represented by the insertions and need to be taken out of the decoy. Therefore, remaining insertions are aligned to the decoy contigs and if there are alignments found, those regions on the decoy are masked with N bases.

**Multimap filter** As more edges are added to the graph, there is a risk of making some regions identical to other parts of the genome, therefore making alignment of reads ambiguous and uninformative. The goal of the multimap filter is to selectively remove graph edges such that the identity of regions is broken. Flow of the process is shown in Figure S1d and details are as follows:

1. Process starts with simulating short reads from all possible haplotypes on the graph. Genome is traversed at specified intervals for start positions and for a given start position, all possible paths of a specified length are generated as reads for that position. All the generated reads are collected as a FASTQ file.
2. Reads generated in the previous step are aligned using the graph
3. Resulting alignments are analyzed for potential bad edge or edge combinations
   (a) Reads at same starting position are grouped
   (b) Within the group there will be one linear reference only read and the rest will be reads following possible combinations of edges on the graph
   (c) Edges used for a non-reference reads is identified and that edge combination is classified as:
      - *BAD*: if read has a mapping quality of zero where reference read had mapping quality greater than zero
      - *GOOD*: if read has a mapping quality greater than the mapping quality of reference read or 20
      - ignored otherwise
   (d) If there are more than one classification of the same edge combination (i.e. different starting position), they are aggregated.

(e) Edge combination is flagged for filtration if it has only *BAD* reads

(f) A minimal subset of edges are identified from the flagged edge combinations, such that each flagged combination will have at least one common edge with the subset

(g) Identified edges within the subset are removed from graph and final graph is outputted

##### S1.2 Allele frequency cutoff experiments

To be able to obtain a more relevant and representative population-specific graph reference, we experimented with different allele frequency (AF) cutoffs to select the common variants to that population. We used Genome-in-a-Bottle (GIAB) Ashkenazim samples HG002, HG003 and HG004 and East Asian samples HG005, HG006 and HG007[1]. We then used gnomAD v3 variants to construct Ashkenazim (N=1,662) and East Asian (N=1,567) graph references at different AF cutoffs (0.5%, 1%, 5%, 10% and 20%)[2]. We ran the Seven Bridges GRAF Workflow for every sample and graph reference constructed to obtain alignment and variant calling results. We also included previous alignment and variant calling results for BWA+GATK and Seven Bridges GRAF using the public Seven Bridges Pan-Genome Graph Reference v1 to see the improvement in using population-specific graph reference compared to linear approach and global graph reference. We calculated comprehensive alignment quality control metrics including graph-specific metrics. We also used SB VCF Benchmark app to benchmark variant calling results against the sample truth sets and Mendelian Violation Detector app to determine the trio concordance among variant calling results.

The alignment QC results show that using graph references increases informative alignment (mapping quality or MAPQ $\geq 20$), reduces multi-mapped (MAPQ$=0$) and uninformative (MAPQ$< 20$) alignments, reduces improperly paired reads and error rate. The use of population-specific graph reference further improves alignment QC. There is an increase in the unmapped reads when we compare BWA+GATK and Pan-Genome; however, some of the increase can be avoided using population-specific graph references as the results show (Figure S3).

The benchmark results show that with the graph references, there is a significant improvement in F-measure, reduction in the total error (the sum of false positives and false negatives) and the variants are more concordant within trios. The use of population-specific graph references further improves these metrics especially in SNP F-measure, SNP total error and INDEL discordance (Figure S4).

The overall results suggest that the use of population-specific graph references is better and the 5% allele frequency cutoff seems working the best for the majority of the metrics and it can be selected as a candidate for the future AF cutoff during the population-specific graph construction.

##### S1.3 Content of the graph references used or constructed

We used the public Seven Bridges Pan-Genome Graph Reference v1 as the global graph reference and we constructed several population-specific graph references in this study[3]. The first population-specific graph was constructed by using only the public resources and the subsequent population graph references were iteratively augmented with the common cohort variants to this graph with the public resources.

The public Seven Bridges Pan-Genome Graph Reference v1 (Pan-Genome) has been constructed using the GRCh38 human reference main contigs as the backbone and GRCh38 human reference alternative contigs, 1000 Genome Project common variants (N=2,504, AF$>$ 1%), Simons Genome Diversity Project common variants (N=278, AC$>$ 10) and the indels discovered by Mills et al (N=36, entire set) and 1000 Genomes consensus indels (N=2, entire set) were augmented to the main contigs in the graph[4–6]. The number of edges each source provides to the Pan-Genome graph is given in Supplementary Table S3.

The population-specific graph with the public resources (Pan-African-0) was constructed using the gnomAD v3 variants that are common in African population (N=21,042, AF_afr$\geq$ 5%). These common

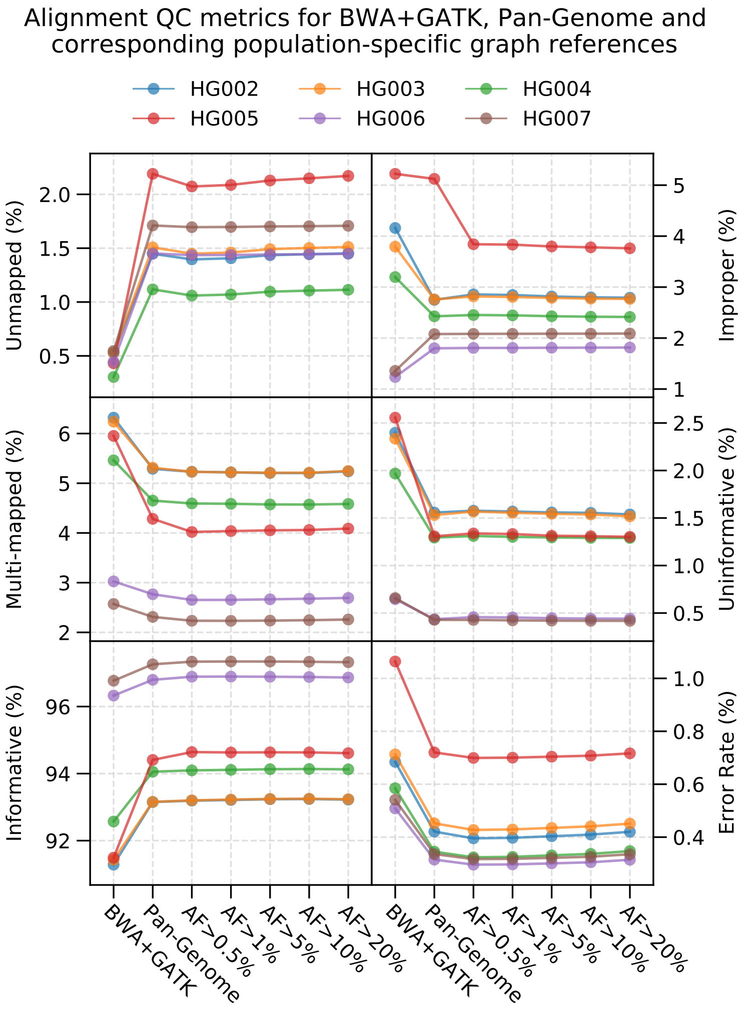


**Figure S3.** **Alignment metrics for GIAB samples.**

African gnomAD v3 variants were augmented with GRCh38 human reference alternative contigs to the GRCh38 human reference main contigs as the backbone [cite gnomAD]. The iterative population-specific graph references (Pan-African-i where i={1, 2, 3, 4, 5}) were constructed using the resulting variant set at each previous group analysis (N=104, AF$\geq$ 5%) in addition to the sources used in the Pan-African-0 graph. List of samples in each group and the benchmarking set is provided in Supplementary Table S14. We also augmented the structural variants (N=10, entire set) discovered by the Human Genome Structural Variation Consortium (HGSVC) in the iterative population-specific graph references[7]. Moreover, we generated a non-iterative Pan-African graph using combined construction variant sets that are subsetted from the variant sets produced by NYGC using the 1000 Genomes high coverage datasets[8]. The number of edges each source provides to the population-specific graph references is given in Supplementary Table S3.

Contents of graph references are shown in Supplementary Table S2. Pan-African graphs include less SNPs than Pan-Genome graph but they contain more insertion and deletions, and difference further increases with addition of African SVs in Pan-African-1 stage. As new groups are included in the graph each type of variant slowly increased. While the Pan-Genome graph includes the more edges than some of the population-specific graphs, the average per sample use rate is lower. Out of 12.5 million SNPs

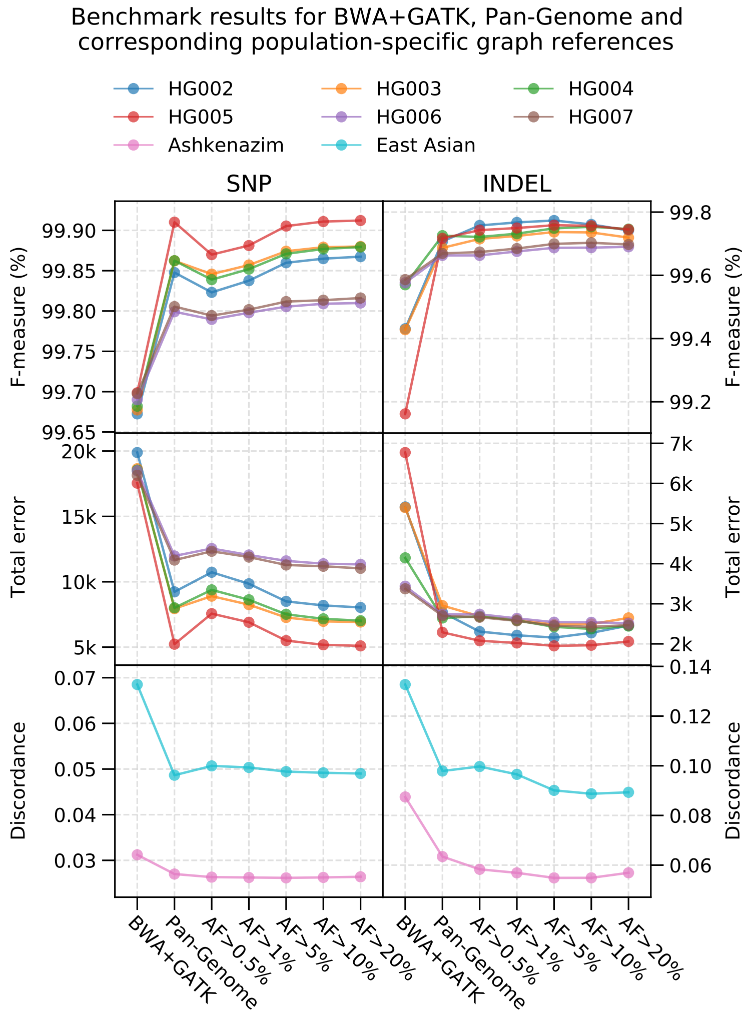


**Figure S4. Benchmark results for GIAB samples.**

only 35% of them are used in the benchmark set for Pan-Genome, but 43% of 12 million SNP edges are used in Pan-African 5. Used deletion edge rate is also 35% for Pan-Genome graph while it is 46% for Pan-African. The most used edge type is insertion, with 77% of Pan-Genome and 80% of Pan-African graph insertions are used in the benchmark set. A non-iterative graph constructed using BWA+GATK results is also compared to the other graphs. It includes the least amount of SNP edges and on average the use edge rate of SNPs is as high as population specific graphs. However, since BWA+GATK graph contains cohort specific INDELs as well as the additional SVs, INDEL use edge rate is high similar to other population-specific graphs.

Finally, we look at the variant composition of the construction sets that are used to construct the graphs *Pan-African 1-5*. The comparison of variants found in each construction set is shown in Figure S5. It is seen that most of the common variants (around 94%) with $AF \geq 5\%$ are shared in all sets as expected, since all sets contain the same number of samples from each African subpopulation and the same female/male ratio. This implies that the processing order of these sets in the project workflow is unimportant. We also observe a significant reduction in the number of variants shared by subsequent groups and the mean AF of the variants added. This shows that most of the more common variations are captured in earlier iterations, and subsequent groups contribute mostly infrequent variants around AF cut-off. The total number of variants for each set is shown in the horizontal bars. A consistent increase is observed with each iteration. This indicates that the variant calling sensitivity increases with each graph augmentation and there is merit in augmenting graphs even during graph constructions.

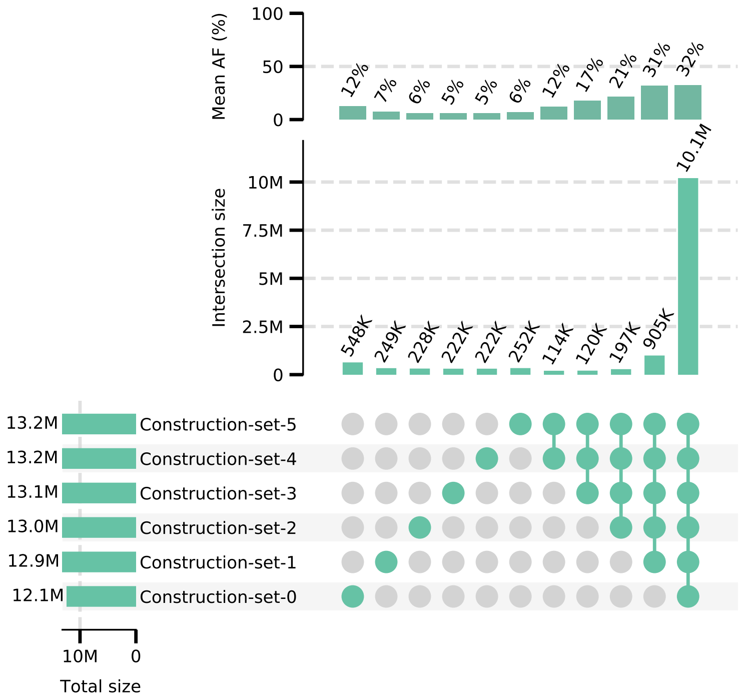


**Figure S5. Comparison of construction set variants.**

#### S2 Nucleotide diversity and absolute divergence

We calculated the nucleotide diversity for all major populations and corresponding subpopulations from the 1000 Genomes dataset. Results are shown in Figure S6. With the exception of American population, subpopulations generally have similar diversity as their populations. America displays a varying diversity among the subpopulations.

The TPR and FPR are also calculated theoretically, assuming an underlying AF distribution obtained empirically for each population, as shown in the inset (see Methods for details). The results are shown in Figure S7a. It is seen that the TPR and FPR improve with increasing number of samples albeit with diminishing returns for larger sample sizes. For instance, by constructing a graph using approximately 700 African samples, one can expect to capture 98% of the variation information with a false positive rate slightly above 2%. The experiment and theory agree well, with some deviation for large number of samples due to the limited number of samples available in the experiment. It is important to note that the rate of improvement with increasing number of samples for each population is inversely correlated with the nucleotide diversity of the population (Figure S6). The descending order of nucleotide diversity is $AFR > AMR > SAS > EUR > EAS$, which is exactly the same order of improvement rates, i.e. convergence rates, from low to high. For instance, we observe a slower convergence for the African population as opposed to the European population because the former is much more diverse. The East Asian population graph, on the other hand, improves the fastest since it has the lowest nucleotide diversity. It is also noteworthy that the convergence rate does not depend on the absolute divergence from GRCh38, as mentioned earlier.

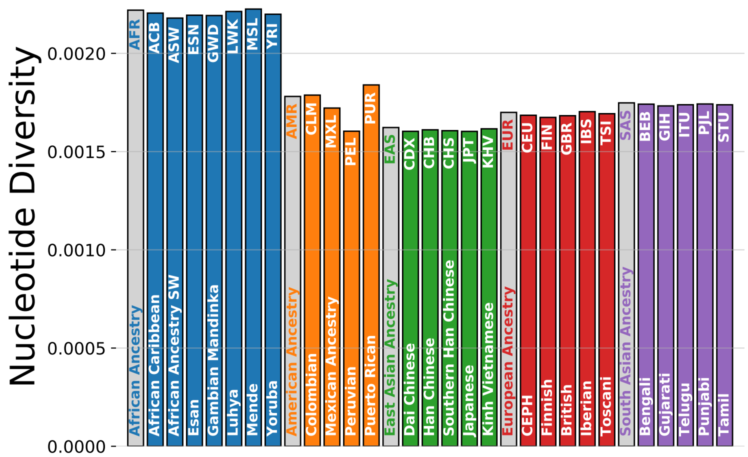


**Figure S6.** **Nucleotide diversity for all subpopulations in the 1000 Genomes dataset**

This is illustrated by the East Asian TPR and FPR curves, which have the highest convergence rates even though the divergence of the East Asian population is higher than all populations except the African. A detailed table for different values of AF cutoff and TPR is provided for all population along with the corresponding FPR in Supplementary Table S1. This table can be used as a guideline for other populations via a comparison of their nucleotide diversity.

In order to measure the effect of sampling on the graph convergence rates, we compare a *homogeneous* sampling approach to a *clustered* sampling approach. Homogeneous sampling ensures that samples for graph construction are picked from all subpopulations uniformly so that there are no abrupt changes in the genetic architecture of the graph reference as more samples are added. Although this approach is expected to more quickly capture the populations genetic information, it is not always possible due to missing detailed ancestry information or different subpopulations being sequenced in different phases of the project. To compare homogeneous sampling to the worst case scenario, clustered approach assumes that the samples are first picked exclusively from a specific subpopulation until there are no samples left belonging to that subpopulation. Then, the procedure is applied to another subpopulation until all subpopulations in the dataset are exhausted. The results are shown in Figure S7b. It is clear that clustered sampling results in a slower convergence rate; therefore, graph references are significantly less representative of the population compared to homogeneous sampling. Abrupt jumps are also observed in TPR and FPR curves at points where a new subpopulation is introduced to the graph reference.

We also calculated TPR and FPR rates for different cutoffs (AF$\geq$ 0.01, AF$\geq$ 0.10 and AF$\geq$ 0.20) in addition to the AF$\geq$ 0.05 shown in the main text. Results are shown in Figure S8. Expected number of samples and corresponding FPR rates are also calculated for various TPR rates and listed in Supplementary Table S1. Number of samples required to reach a target TPR follows the diversity for correspoding populations. The more diverse the population is, the more samples are required to get a representative graph.

### S3 Alignment Statistics

#### S3.1 Alignment QC

We determined the alignment quality of reads through commonly used metrics like the number of mapped reads, proper reads, mapping quality, error rate and coverage. We compared population specific Pan-

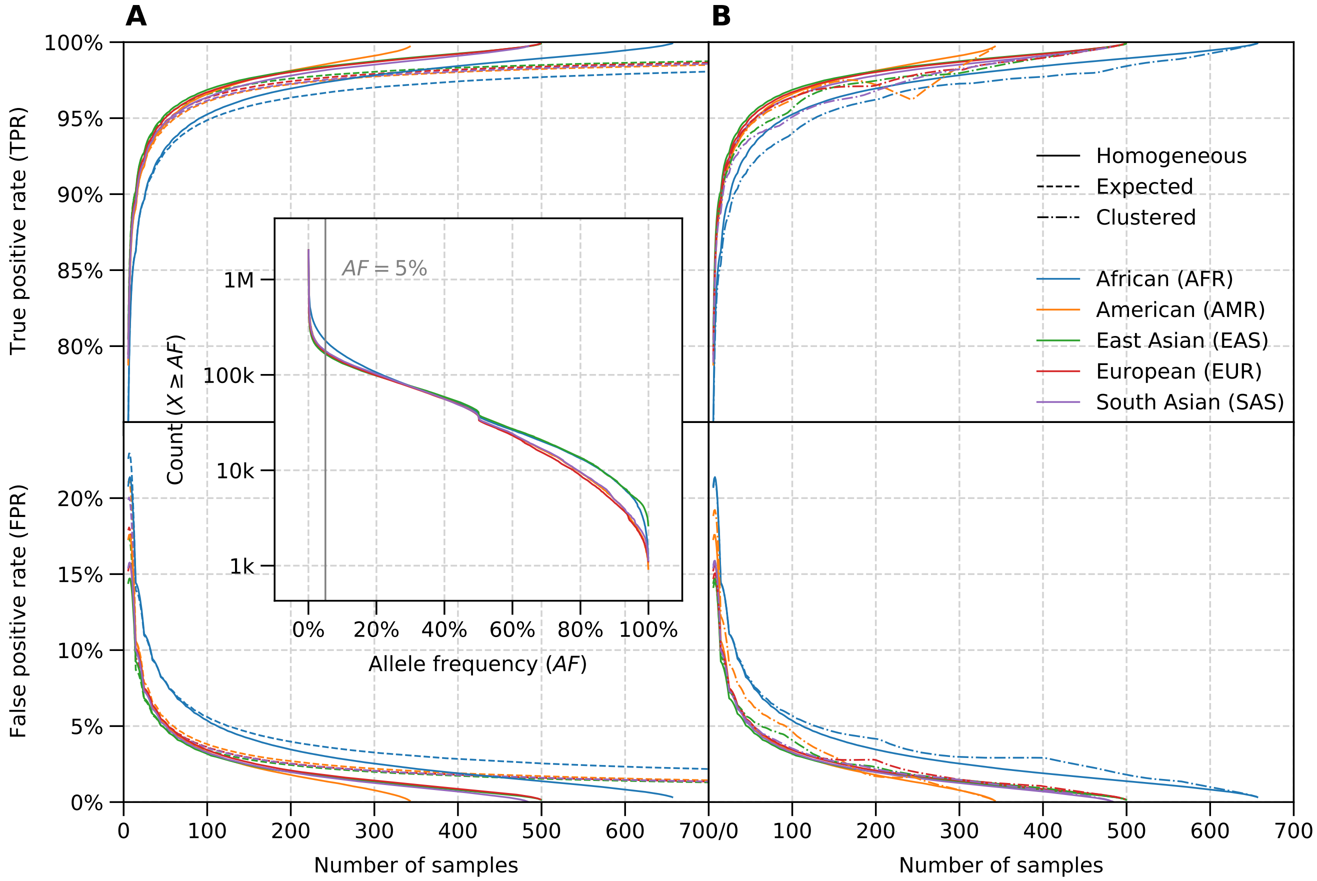


**Figure S7.** **True positive rate (TPR) and false positive rate (FPR) in population-specific graph references with respect to number of samples used for graph construction.** TPR and FPR are calculated for each population shown in the legend. An allele frequency (AF) threshold of 5% is used, below which the variants are discarded. (a) Theoretical calculation (see Methods) and simulated results. (b) Simulated results with homogeneous and clustered sampling. Homogeneous sampling assumes samples are taken from subpopulations uniformly, whereas clustered approach simulates the extreme case where samples are taken from each subpopulation until it is exhausted. Inset shows the AF distribution used for calculations, which is obtained empirically from the 1000 Genomes dataset for each major population.

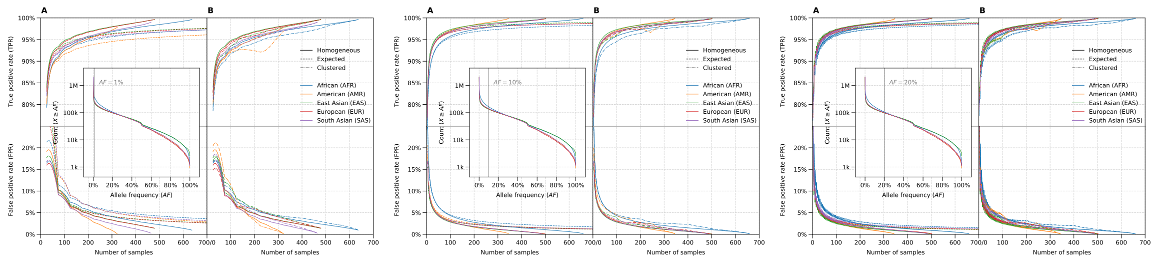

**Figure S8.** **TPR and FPR rates for different AF cutoffs (0.01, 0.1 and 0.2) for populations in 1000 Genomes data set**

African graphs to Pan-Genome graph and linear reference based BWA+GATK workflow. Means of metrics over all benchmark results are presented in Supplementary Table S8. We observed that BWA reported more aligned reads, consistent with the more lenient approach. Graph workflows resulted with less improper reads and error rates. Decrease in improper reads is by nearly 5 million and error rate decreased by nearly half for the final population-specific graph. Also, we see that while BWA+GATK mapped on average 1.1 million more higher confidence reads (MAPQ$\geq$ 20), it includes 2.4 million more multi-mapped reads (MAPQ$=$ 0) and 5.5 million more low quality alignments (MAPQ$<$ 20). Additionally, graph generated directly from BWA+GATK variants instead of an iterative approach also shows a smaller range of error rate, improper reads and unmapped reads as well as less multi-maped and low quality reads and more high quality reads than linear BWA+GATK.

Population-specific graphs improved over Pan-Genome results. For instance, the number of mapped reads, high confidence reads and mean coverage increased in Pan-African iterations. Also, improper reads, low quality reads and error rate further decreased in Pan-African graph results. The number of multi-mapped reads initially increased in the first Pan-African graph but later graphs showed the decreasing trend.

##### S3.2 Read Relocation Between Pipelines

Alignment results of the benchmark sample HG02470 are compared between BWA+GATK, Pan-Genome and Pan-African-5 workflows in order to get an insight on the quality of the reads that change locations. Reads that map to different locations are identified and analysed. Results are presented in Supplementary Table S9.

In the first table, reads that are aligned to the same chromosome with different positions are compared. Most notable difference is when the read alignment confidence changes. Graph was able to identify more alignments with higher confidence (MAPQ $\geq$ 20) where BWA aligned with lower confidence (MAPQ $<$ 20). In the second table, similar comparison is performed but for read that map to different chromosomes or left unmapped. Graph pipeline results in more unmapped reads compared to BWA. This is due to the lenient approach used by BWA compared to graph aligner for reporting read alignments by default. Another major difference is the number of low and high confidence reads. Graph aligner produces more high confidence reads compared to BWA and population-specific graphs further increases this compared to Pan-Genome

#### S4 Variant Statistics

##### S4.1 Variant QC

Variants grouped by types are analyzed and shown in Supplementary Table S4. Graph pipelines called more variants compared to BWA+GATK. There is a 10.2% increase in average total variant counts for Pan-African 5 graph. Population specific Pan-African graph resulted with additional 450,000 known SNPs and 6,041 known INDELs (with dbSNP v154) and additional 28,900 novel SNPs and 73,553 novel INDELs compared to BWA+GATK.

In order to understand the quality of these incoming variants, we analysed Het/Hom and Ti/Tv ratios which are well known to be population specific metrics for evolutionary reasons[9]. Het/Hom rate of known variants is around the expected 2.0 for all workflows Graph novel SNP Het/Hom rates are higher than BWA+GATK and using Pan-African graphs lead to a small decrease yet we observed an increase in novel INDEL Het/Hom rates using Pan-African graph and the rate is higher than BWA+GATK. Ti/Tv rate appears to be very stable in the workflows, a small difference is observed in graph results, novel Ti/Tv rates are higher in graph results and known Ti/Tv rate is slightly higher in BWA+GATK. Thus, both

Het/Hom rate and ts/tv rate is quite similar in workflows indicating the quality of the variants is high. The non-iterative graph generated using BWA+GATK variants display a lower rate of Ti/Tv than linear BWA+GATK and higher rates of Het/Hom for all of the metrics separately.

Considering the total number of variants of samples from merged VCFs, Pan-African benchmarks, similar to per sample variants, include more SNPs and INDELs than BWA+GATK, the last Pan-African resulted with 2.4 million more known and 606 thousand novel SNPs and 187 thousand more known and 1.9 million more novel INDELs than BWA+GATK. Singleton rates are approximately same, nearly 30% in Pan-African and 29.4% in BWA+GATK of known SNPs is singleton and 29% in Pan-African and 23% in BWA+GATK of known INDELs is singleton. On the contrary, while the novel numbers are smaller than knowns, we observed that 67% in Pan-African and 79.7% in BWA+GATK of novel SNPs is singleton. Novel INDEL singleton rates are smaller than SNPs, 27% in Pan-African and 15.6% in BWA+GATK of novel INDELs is singleton. The graph generated using BWA+GATK graph includes more known but less novel SNPs and INDELs than Pan-African iteration graphs yet it includes more known but less novel singletons.

Based on the AF distributions, graph generated higher number of unique and common variants in higher frequencies. Furthermore, we noticed that, the number of known SNPs and INDELs decreases as the graph becomes more population specific comparing Pan-Genome to Pan-African graphs, and the number of novel SNPs and INDELs increases while the rate of singletons not changing too much. Ti/Tv rates for graph benchmarks are very generally similar but they are smaller than BWA+GATK Ti/Tv rates.

The number of total SNPs and INDELs for population specific Pan-African graphs, Pan-Genome graph and BWA+GATK analysis are compared in difficult regions provided by Genome in a Bottle[10] and shown at Supplementary Table S13. The variants are mapped to the regions in two aspects: one as difficult regions vs the rest and second as low mapping and segmental duplications vs rest. We observed that in all regions, the graph pipeline is able to find more SNPs and more INDELs without any bias to any region. Both Pan-Genome and Pan-African graphs able to call more SNPs and INDELs than BWA+GATK in both difficult regions and low mapping and segmental duplication regions.

Original variant counts were shown without any filtration. In order to understand the effect of basic VCF filtering based on variant quality to the number of total variant calls with known vs novel distinctions, Het/Hom rates and Ti/Tv ratios, per sample VCF files were filtered with different QUAL (¿30, ¿50 and ¿70) values. Only the final graph (Pan-African 5) analyzed and compared to non-filtered metrics in this step. Details of this analysis can be found in Supplementary Table S12. We observed that the number of filtered variants is between 33,180 and 115,995 which is a small quantity of the total variant set (6,226,870). Yet, in filtered set of variants Het/Hom rate is decreased between 0.02 to 0.2 points and Ti/Tv rate is increased by 0.02 points. Furthermore, in merged set of variants the effect is sharper; we saw that the Ti/Tv ratio can be increased by 0.03 points for novel variants by filtering off only 298,706 variants.

#### S4.2 Joint calling only regions

GATK team recommends joint analysis on a set of samples to perform variant discovery in their Best Practices workflow in order to empower population-wide information increasing sensitivity and accuracy over calling single samples. Since population specific graphs found more frequent variants in the African population compared to GATK joint call results, we investigated whether graph can call the regions that are uniquely called by joint calling but not in single sample calls where they suggest population specific information is enriched.

We compared joint call results of the benchmark samples with GATK's single call results and counted those genotypes in two respects; homozygous (1/1 in joint call vs no call in single call) and heterozygous (0/1 in joint call vs no call in single call). Results are shown in Supplementary Table S6. We found that joint calling mostly rescues 0/1 genotypes (22,424,588 for 0/1 vs 1,713,068 for 1/1) and from these heterozygous genotypes, INDEL variant type is more than SNPs suggesting that single variant calling mostly loses heterozygous INDELs. Furthermore, most of these variants were tagged as PASS by VQSR. Conversely, most of the heterozygous SNPs rescued by joint calling (56,397) is tagged as non-PASS. Therefore, joint-calling is advantageous over single calling in heterozygous INDEL calling but not SNPs.

In our analysis, we hypothesized that population specific graphs already include relevant variant information and could potentially identify these rescued sites. We found out that population specific graphs can find most of the genotypes that were rescued by joint calling in samples. Population specific graphs are able to find about 77% of 13,626,564 0/1 PASS genotypes and 86% of 1,461,641 1/1 PASS genotypes. In addition to that, we observed that most of the genotype missed using single call is INDEL type, and population specific graphs able to identify 83% of 12,934,650 PASS INDELs while calling only 28% of 902,537 non-PASS INDELs. Out of 2,153,555 PASS SNPs, population specific graphs successfully called 47% of them while calling only 17% of 8,146,914 non-PASS SNP regions. Finally, we confidently can say that population specific graphs are more likely to identify joint-only regions that passed the VQSR filter. Pan-African graphs found nearly 78% of 15,088,205 PASS tagged regions while identifying only 18% of 9,049,451.

##### S4.3 Allele frequency distribution

We counted the number of SNPs and INDELs detected by GRAPH (with Pan-African-5) and the GATK and binned the counts by allele frequency (AF) in the bin size of 1% to investigate the sensitivity of the two pipelines. We counted the variant that are commonly and uniquely detected in both pipelines. We also divided commonly detected variants into to by the AF difference where in the first category the variants have higher AFs in GRAPH ($AF_{GRAF} > AF_{GATK}$) and in the second category the variants have higher AFs in GATK ($AF_{GATK} > AF_{GRAF}$). The cumulative distribution of these counts is given in Figure 14. The counts for this analysis are given in Supplementary Table S7.

##### S4.4 Functional effect analysis

In order to evaluate the functional importance of the discovered variants, we used Ensembl Variant Effect Predictor[11] (VEP) GRCh38 release 101 on the unique GRAPH results with the Pan-African-5 population graph reference and the unique GATK results that are generated by NYGC for the benchmark samples (N= 141). We used the `--flag_pick` option in VEP to select the annotation per transcript and used the picked annotation. We stratified the VEP results based on the IMPACT score defined by VEP (HIGH, MODERATE, LOW and MODIFIER), genomic region using the annotation definitions (exonic, intronic, intergenic) and allele frequency (AF) categories (Singleton where the variant is discovered in only single individual as heterozygous or homozygous, Rare where the variant is not a Singleton and has AF$<$ 5%, Common where the variant has AF$\geq$ 5%). The breakdown of the consequences of the VEP annotation is given in Supplementary Table S5.

#### S5 Computational Efficiency Measurements

Execution details are summarized in the table 1. Graph and index construction time denotes the time it takes aligner to create graph along with search index from the input FASTA and VCF file. There is a slight correlation with the number of variants in the graph but total time is considerably fast (below 3 minutes). Graph and index memory usage is the aligner memory usage right after graph and index is constructed and before alignment starts. It is consistently around 16GB with minor variations due to the variants included in the graph. Alignment time shows the mean duration for alignment of benchmark samples. Alignment time is affected by the number of variants as well as the composition of added variants. The jump at Pan-African 1 in alignment time mostly due to the addition of larger structural variants, specifically the insertions. Variant calling time follows the similar trend in alignment time, however the jump at Pan-African 1 is considerably smaller.

| Graph name | Graph and index construction time | Graph and index memory usage | Alignment time | Variant call time |
|---|---|---|---|---|
| PanGenome | 164s | 16.70 GB | 2h 42m | 41m |
| Pan-African 0 | 154s | 16.01 GB | 2h 54m | 41m |
| Pan-African 1 | 164s | 16.43 GB | 4h 21m | 46m |
| Pan-African 2 | 164s | 16.71 GB | 4h 31m | 47m |
| Pan-African 3 | 173s | 16.69 GB | 4h 38m | 47m |
| Pan-African 4 | 173s | 16.73 GB | 4h 41m | 47m |
| Pan-African 5 | 174s | 16.85 GB | 4h 43m | 47m |

**Table 1.** Execution details for graph, alignment and variant calling

### S6 Bioinformatics pipelines and command lines

#### S6.1 Seven Bridges Germline Pipeline

***Input Files***

- CRAM reference: GRCh38_full_analysis_set_plus_decoy_hla.fa
- CRAM reference index: GRCh38_full_analysis_set_plus_decoy_hla.fa.fai
- CRAM reference dictionary: GRCh38_full_analysis_set_plus_decoy_hla.dict
- Intervals: GRCh38.GRAF.Genome_Intervals.v1.bed
- Graph reference: GRCh38.GRAF.Reference.vcf.gz
- Graph reference index: GRCh38.GRAF.Reference.vcf.gz.tbi
- Linear reference: GRCh38.Linear.GRAF.Reference.fa
- Linear reference index: GRCh38.Linear.GRAF.Reference.fa.fai
- Input CRAM file: sample_name.cram
- Input CRAM file index: sample_name.cram.crai

***Output Files***

- Alignments: sample_name.bam
- Filtered variants: sample_name_filtered.vcf
- Variant stats: sample_name.stats
- Graph alignment summary: sample_name-summary.json
- Graph used edge data: sample_name.json.gz

##### ***Workflow Steps and Tools***

- Read Processing: Samtools 1.9

```
samtools collate -u -O --threads 31 -n 64 --reference
  GRCh38_full_analysis_set_plus_decoy_hla.fa
  sample_name.cram |
  samtools view -F 2304 -o sample_name_collated.cram
  --reference GRCh38_full_analysis_set_plus_decoy_hla.fa
  --output-fmt CRAM --threads 31 -
```

- Alignment: Seven Bridges Graph Aligner (rasm) 1.0rc3

```
aligner --cram_ref GRCh38_full_analysis_set_plus_decoy_hla.fa
  --vcf GRCh38.GRAF.Reference.vcf.gz
  --reference GRCh38.Linear.GRAF.Reference.fa
  --markdup --fmt bam
  -q sample_name_collated.cram
  --read_group_platform 'Illumina'
  --read_group_sample sample_name
  --read_group_library sample_name
  --sort --tmp ./sort.chunk.XXXXXX
  --sort_mem 30000 --hts_threads 40
  --merge_threads 12 --keep
  -o sample_name.bam &&
  sambamba index -t 36 sample_name.bam
```

- Variant Calling: Seven Bridges Reassembly Variant Caller 1.0rc3

```
rasm -v sample_name.vcf -a GRCh38.GRAF.Genome_Intervals.v1.bed
  -f GRCh38.Linear.GRAF.Reference.fa -b sample_name.bam
  -g GRCh38.GRAF.Reference.vcf.gz -x all -s 10 -t 0
```

- Filtering: Bcftools 1.9

```
bcftools filter -o sample_name_filtered.vcf
  -e '(TYPE="snp" && (INFO/AD_Ratio[1] < 0.20 ||
  INFO/MBQ[1] < 15 || INFO/QD < 1 ||
  INFO/MQRankSum < -8 || INFO/FS > 50)) ||
  (TYPE="indel" && INFO/AD_Ratio[1] < 0.15)'
  -O v -s FP -m x sample_name.vcf
```

- Alignment and Variant stats: Graph Stats 1.0 & Bcftools 1.9

```
python graph_stats.py --bam sample_name.bam
  --vcf GRCh38.GRAF.Reference.vcf.gz --pretty
```

```
bash gzip_vcf.sh &&
  bcftools stats
  --fasta-ref GRCh38.Linear.GRAF.Reference.fa
  sample_name.vcf.gz > sample_name.stats;
  mkdir plots_dir &&
  plot-vcfstats sample_name.stats -p plots_dir/
```

##### S6.2 Collect Benchmark Stats Pipeline

###### *Input Files*

- Annotation file: HG38.dbsnp154.vcf.gz
- Annotation file index: HG38.dbsnp154.vcf.gz.tbi
- Ploidy definition file:ploidy_hg38_vcffix.txt
- Reference sequence in FASTA format: GRCh38.Linear.GRAF.Reference.fa
- Reference sequence in FASTA format index: GRCh38.Linear.GRAF.Reference.fa.fai
- Samples list: sample_genders.txt
- Input variants file array: sample_name_1.vcf, sample_name_2.vcf ..

###### *Output Files*

- Filtered output VCF file: benchmark.vcf.gz
- Bcftools stat output file: benchmark.stats
- RTG vcfstats output file - known: benchmark.known.vcfstats
- RTG vcfstats output file - novel: benchmark.novel.vcfstats
- Counts: benchmark.frq.count
- Frequency: benchmark.frq
- HWE: benchmark.hwe
- Heterozygosity: benchmark.het

###### *Workflow Steps and Tools*

- Normalization: Bcftools 1.9

```
bgzip -c -f sample_name.vcf > sample_name.vcf.gz &&
  bcftools index -f -t sample_name.vcf.gz &&
  bcftools norm --threads 16
  --fasta-ref GRCh38.Linear.GRAF.Reference.fa
  --multiallelics -any --output sample_name.norm.vcf.gz
  --output-type z sample_name.vcf.gz
```

- Merging: Bcftools 1.9

```
bash gzip_vcf.sh &&
  bcftools merge  --output benchmark.merged.vcf.gz
  --threads 16 --missing-to-ref --merge none
  --output-type z
  [sample_name_1.norm.vcf.gz  sample_name_2.norm.vcf.gz ..]
```

- Annotations and Ploidy Fix: Bcftools 1.9 & VCFtools 0.1.14

```
bcftools index  -f -t benchmark.merged.vcf.gz &&
  bcftools annotate --output benchmark.merged.annotated.vcf
  --set-id . --remove "INFO" benchmark.merged.vcf.gz

outpbgzip -c -f benchmark.merged.annotated.vcf
  > benchmark.merged.annotated.vcf.gz &&
  bcftools index -f -t benchmark.merged.annotated.vcf.gz &&
  bcftools annotate --output benchmark.annotated.vcf
  --annotations HG38.dbsnp154.vcf.gz --columns ID
  --output-type v --remove "FILTER" --threads 16
  benchmark.merged.annotated.vcf.gz

cat benchmark.merged.annotated.vcf | vcf-fix-ploidy
  --fix-likelihoods --ploidy ploidy_hg38_vcffix.txt
  --samples sample_genders.txt > benchmark.annotated_fixed.vcf

bcftools +fill-tags -o benchmark.annotated_fixed.info.vcf
  -Ov benchmark.annotated_fixed.vcf
```

- Filtration: Bcftools 1.9

```
bgzip -c -f benchmark.annotated_fixed.info.vcf
  > benchmark.annotated_fixed.info.vcf.gz &&
  bcftools index -f -t benchmark.annotated_fixed.info.vcf.gz &&
  bcftools filter --output benchmark.vcf.gz --include 'AC>0'
  --output-type z --threads 16
  benchmark.annotated_fixed.info.vcf.gz
```

- Variant Statistics: Bcftools 1.9 & VCFtools 0.1.14 & RTG Tools 3.6.2

```
bash gzip_vcf.sh &&
  bcftools stats --split-by-ID --samples -
  --fasta-ref GRCh38.Linear.GRAF.Reference.fa
  benchmark.vcf.gz
  > benchmark.vcf.stats; mkdir plots_dir &&
  plot-vcfstats benchmark.stats -p plots_dir/

rtg vcfstats --novel benchmark.vcf.gz
```

```
  > benchmark.novel.vcfstats
rtg vcfstats --known benchmark.vcf.gz
  > benchmark.known.vcfstats

vcftools --counts2 --gzvcf benchmark.vcf.gz
  --out benchmark.frq.count
vcftools --het --gzvcf benchmark.vcf.gz
  --out benchmark.het
vcftools --hardy --gzvcf benchmark.vcf.gz
  --out benchmark.hwe
vcftools --freq2 --gzvcf benchmark.vcf.gz
  --out benchmark.frq
```
